# Efficient synthesis of arabinosyl nucleosides via active site engineering of the *Leishmania mexicana* purine 2′-deoxyribosyltranferase

**DOI:** 10.1101/2024.11.29.626018

**Authors:** Davide A. Cecchini, Javier Acosta, Laura P. Saiz-Álvarez, Carmen Ortega, Pierre Alexandre Kaminski, Jesús Fernández-Lucas, Aurelio Hidalgo

**Affiliations:** Centro de Biología Molecular Severo Ochoa (UAM-CSIC), Nicolás Cabrera 1, 28049 Madrid, Spain; Department of Molecular Biology, Universidad Autónoma de Madrid, Campus de Cantoblanco, 28049 Madrid, Spain; Applied Biotechnology Group, Universidad Europea de Madrid, Urbanización El Bosque, 28670 Villaviciosa de Odón (Madrid), Spain; Institut Pasteur, Université Paris Cité, Unité Plasticité du Génome Bactérien, CNRS UMR3525, 75724 Paris, France; Department of Biochemistry and Molecular Biology, Faculty of Biology, Universidad Complutense de Madrid, C. de José Antonio Novais, 12, 28040 Madrid, Spain; Grupo de Investigación en Ciencias Naturales y Exactas, GICNEX, Universidad de la Costa, CUC, Calle 58 # 55 – 66, 080002 Barranquilla, Colombia; Instituto Universitario de Biología Molecular (IUBM), Universidad Autónoma de Madrid, Campus de Cantoblanco, 28049 Madrid, Spain

**Keywords:** protein engineering, biocatalysis, structural bioinformatics, nucleoside analogs

## Abstract

Even though 2′-deoxyribosyltransferases (NDTs) have emerged as promising and eco-friendly catalysts for synthesizing different 2’-deoxynucleoside analogs, their limited activity towards substrates featuring modifications at the 2′ -C and 3′ -C positions of the ribose moiety significantly restrict their potential application in the pharmaceutical industry. Herein we report two engineered variants from *Leishmania mexicana* purine deoxyribosyltransferase (*Lm*PDT) with the highest reported arabinosyltransferase activity for an NDT, achieved through a semi-rational design approach. The resulting variants, *Lm*PDTN56L and *Lm*PDTN56C, display up to 13.5- and 30-fold enhanced activity on the synthesis of vidarabine compared to the wild-type enzyme. Moreover, thermal unfolding and thermal challenge experiments revealed that the improvement in arabinosyl transferase activity did not result in a stability trade-off, with *Tm* values similar to the wild-type enzyme and similar activity retention at 40 °C. After biochemical characterization, the optimal reaction parameters for variant *Lm*PDTN56C were determined to be pH 6 and a temperature interval from 30 to 45 °C. Interestingly, under optimal operational conditions (40 °C, 50 mM MES buffer pH 6) the initial 30-fold enhancement observed for *Lm*PDTN56C escalated to a remarkable 40-fold improvement. This comprehensive study elucidates the potential of the engineered *Lm*PDT variants for efficient and tailored biocatalytic synthesis of arabinosyl nucleosides, paving the way for enhanced applications in nucleoside analog production.

## INTRODUCTION

Nucleoside analogs (NAs) are significant synthetic targets in medicinal chemistry, primarily as inhibitors or precursors thereof in the treatment of cancer and viral diseases,^[1,2]^, but also as noncanonical building blocks for synthetic RNA and DNA oligonucleotides^[3]^. Their action relies on their similarity to biomolecules involved in DNA and RNA synthesis or cell signalling, diverging from the natural nucleosides in the sugar moiety (e.g. vidarabine, didanosine, the nucleobase (e.g. ribavirin) or both (e.g. islatravir, lamivudine, emtricitabine, abacavir). In particular, nucleosides, in which the 2’deoxyribose moiety has been replaced with arabinose, or arabinosyl nucleosides have found application as antiviral or antineoplasic. For instance, arabinosyl adenine (commonly named Vidarabine), originally intended as an anti-cancer drug, was the first nucleoside antiviral drug to receive approval from the FDA^[1]^ and due to its notable efficacy in addressing infections induced by the herpes simplex virus, and some of its specific clinical manifestations, including herpes simplex encephalitis and herpetic keratitis.

With the expanding relevance of nucleosides in pharmaceutical applications and a growing concern for the efficiency and environmental sustainability of industrial processes as well as a social demand for affordable drugs, there is a growing need for more efficient, safe and sustainable synthetic methods avoiding groups of chemicals that may be harmful to the environment and persistently toxic, acompatible with the bioeconomy and in line with the “safe and sustainable by design” framework^[4,5]^. In this context, enzyme-catalyzed processes emerge as an efficient and eco-friendlier alternative to traditional chemical methodologies for the synthesis of nucleoside analogs^[6–11]^. The transglycosylation reaction catalyzed by nucleoside phosphorylases (NPs)^[12–17]^ and nucleoside-2′-deoxyribosyltransferases (NDTs)^[18–22]^ offer a one-pot, regio-, chemo- and enantioselective synthetic methodology to synthesize nucleoside APIs. This alternative circumvents the challenges associated with chemical processes, such as the need for protecting and de-protecting functional groups, laborious purification of intermediates, and the use of organic reagents and potentially hazardous solvents^[23–26]^.

Despite the significant potential of NDTs as catalysts for the synthesis of nucleoside analogues, their narrow specificity towards sugar-moiety recognition and their limited stability under harsh industrial conditions hinder their applicability in industrial processes. While research labs have invested considerable effort to improve the durability of NDTs through enzyme immobilization^[27–29]^, enzyme engineering^[30]^, or the use of extremozymes^[31,32]^, considerably fewer attempts have been reported towards broadening the substrate scope of NDTs^[33–35]^. However, recent studies have uncovered a number of NDTs with limited activity towards substrates bearing modifications in the 2′-C and 3′-C positions of the 2′-deoxyribose moiety, among them those from *Lactobacilli, Trypanosoma or Archaeoglobus* ^[7,29,36,37]^. Unfortunately, this low-level activity is insufficient to consider these enzymes as industrial biocatalysts for the synthesis of 2′- and 3′-modified nucleosides but constitutes an excellent springboard to explore sequence space for improved fitness by means of protein engineering^[38]^.

The rational prediction of improvements in protein fitness is a complex task because methods are limited by the quality of available sequence-structure-function hypotheses and direct improvements may not always be obtained. Complementarily, directed evolution is independent of structure-function knowledge, at the cost of increasing exponentially the size of the library to be screened to cover all possible combinations of the protein sequence or even a significant amount of sequence space. Hybrid approaches, often termed either semi-rational design or focused directed evolution^[39]^, leverage the best features of each, by narrowing down the areas to be randomized and thus using either sequence and/or structural information to limit the amount of sequence space to be screened^[40]^. Whereas the exploration of targeted sites can be simultaneous, such as in DNA shuffling^[41]^, a stepwise exploration of sites increases the odds for cooperativity and reduces the chances for deleterious or compensatory interactions among the introduced amino acid replacements, leading to high quality libraries^[42–44]^.

Using the latter methodology, we engineered the *Leishmania mexicana* purine deoxyribosyltransferase (*Lm*PDT) for highly efficient production of arabinosyl nucleosides through a semi-rational design approach. This effort yielded two *Lm*PDT variants, *Lm*PDTN56L and *Lm*PDTN56C, with significantly improved arabinosyl transferase activity compared to the wild-type enzyme. Notably, *Lm*PDTN56L exhibited a 13.5-fold increase, while *Lm*PDTN56C demonstrated an even more remarkable 30-fold enhancement in the synthesis of vidarabine. Interestingly, optimizing the operational conditions for using *Lm*PDTN56C led to a 40-fold improvement in the initial arabinosyl transferase activity. Finally, different operational variables were examined including pH, temperature, enzyme stability, catalyst loading and substrate concentration.

## MATERIALS AND METHODS

### Materials

Cell culture medium reagents were procured from Difco (St. Louis, USA), while Merck (Darmstadt, Germany) supplied the triethyl ammonium acetate buffer and biological reagents. All remaining chemical reagents and organic solvents were obtained from Symta (Madrid, Spain). The nucleosides, nucleotides, and nucleobases used were sourced from Carbosynth Ltd. (Compton, UK).

### Strains, constructs, primers and growth media

*E. coli* PAK-6 Δ*guaBguaA*::gm Δdeo-11 was used for library transformation by electrocompetence,^[45,46]^ and selection. *E. coli DH5α* [*supE44, ΔlacU169 (φ80 lacZΔM15), hsdR17, recA, endA1, gyrA96, thi-1 relA1*] was used for construction of plasmids and protein expression. Luria-Bertani (LB) lysogeny broth (10 g/L tryptone, 5 g/L yeast extract, 5 g/L NaCl) was used for general purposes and M9 minimal medium supplemented with 0.2% (v/v) glycerol as carbon source was used for library selection. Both media were supplemented with 100 mg/L ampicillin (Amp), 10 mg/L gentamycin (Gen) and/or 20 g/l agar when required.

Constructs used in this work are detailed in Supplementary Table 1 and oligonucleotides are specified in Supplementary Table 2.

### Library construction and selection of *Lm*PDT variants

Wild-type *Lm*PDT was cloned with a ribosome-binding site and six-histidine tag (His6 tag) at the N-terminus in plasmid pBluescript-SK between the *Xba*I and *Eco*RI restriction sites. Single-site saturation mutagenesis was performed with the Quikchange Site-directed mutagenesis kit (Agilent) on positions Tyr8, Phe15, Glu43, Asn 56, Asp79, Thr82, Glu121, Asn128, Met130 using NNK degenerate codons.

Electrocompetent *E. coli* PAK6 cells were transformed with the generated libraries and plated on M9 medium supplemented with ampicillin, gentamycin, glycerol, 0.3 mM arabinosyl guanine (ara G), 0.3 mM adenine (Ade), and 0.5mM IPTG as inducer. An aliquot was plated on LBAmp plates to estimate the library size and another one on M). Controls with the wild-type enzyme were inoculated on selection plates to determine the background arabinosyl transferase activity of the parental enzyme. Colonies grown after 3-4 days were considered putative hits, replica streaked on LBAmp plates and grown in 5 mL LB medium supplemented with ampicillin. Plasmids were extracted using the Promega Wizard SV Miniprep Kit (Madison, WI, USA) and sent for Sanger sequencing (Macrogen Inc., Seoul, Republic of Korea).

To discard possible effects due to phenotypic variability of the host used for selection, plasmids were retransformed in *E. coli* DH5α cells before verification of the enhanced arabinosyltransferase activity with crude extracts.

### Protein expression and purification

One colony was grown in 5 mL LB medium supplemented with ampicillin for 16 h at 37 °C and 180 rpm. Later, the culture was diluted to OD600 0.05 in 50 mL fresh medium in 250 mL Erlenmenyer flasks. Cultures were incubated at 37 °C and 180 rpm until OD600 0.6, when IPTG was added to a final concentration of 0.5mM. Absorbance was measured at 600 nm in a FLUOstar Optima microplate reader (BMG Labtech GmbH, Ortenberg, Germany). Cell pellets were collected by centrifugation at 4,000 ×*g* and 4 °C for 15 min (5804R centrifuge, Eppendorf), washed with 50 mM Tris-HCl buffer, pH 7.5 and stored at −20 °C until use.

To verify the subcellular location of recombinant proteins, pellets were resuspended in 50 mM phosphate buffer, sonicated (0.6 Amplitude and 50% pulse) and centrifuged at 14,100 ×*g* for 10 minutes. The supernatant was separated and the pellet was washed with phosphate buffer 50 mM pH 7.5 and 0.1% w/v Triton X-100, analyzed by SDS-PAGE in a 12% acrylamide gel according to the method described by Laemmli^[47]^. The gels were stained with Coomassie Brilliant Blue G-250 from Bio-Rad Laboratories (Hercules, USA).

Proteins found in the soluble fraction were later purified by immobilized metal ion affinity chromatography (IMAC), using Talon resin (BD ClonTech). To this end, cell pellets were homogenized (GEA Lab Homogenizer PandaPLUS 2000, GEA Niro Soavi), centrifuged 12,000 ×*g* and 4 °C for 30 min. Cell lysate was applied to the IMAC resin and purified according to the manufacturer’s instructions. Subsequently, proteins were dialyzed against 20 mM Tris-HCl pH7.5 for 16h at 4 C to remove the imidazole, concentrated using 3 kDa cutoff Amicon Ultra Centrifugal Filters (Merck Millipore). Fractions were then applied to a HiPrep 16/60 Sephacryl S-100 gel filtration column using an Akta Start chromatography system (GE Biosciences) using 0.5 ml/min of 50 mM Tris-HCl pH 7.5 as mobile phase and stored at 4 °C until further use. After protein concentration, purity was checked by sodium dodecyl sulfate polyacrylamide gel electrophoresis (SDS-PAGE) in a 12% polyacrylamide gel and their concentration quantitated using the Bio-Rad Protein Assay (Bio-Rad, Hercules, CA, USA), according to the manufacturer’s protocol, using bovine serum albumin (BSA) as standard.

### Glycosyltransferase activity assays

The standard 2′-deoxyribosyltransferase activity assay was conducted by incubating 0.3 µg of pure enzyme with a 40 µL solution containing 10 mM 2′-deoxyinosine (dIno) and 10 mMAde in 50 mM MES pH 6.5 at 40 °C and 300 r.p.m. for 5 min. Subsequently, the enzyme was inactivated, following the procedure previously described^[48]^, and the production of 2’-deoxyadenosine (dAdo) was analyzed and quantified using HPLC. Each determination was performed in triplicate, with a maximum error of less than 5%. Under these conditions, one activity unit (U) was defined as the amount of enzyme (mg) capable of producing 1 μmol/min (IU) of dAdo under the assay conditions.

The standard protocol for assessing arabinosyl transferase activity involved incubating 4 µg of pure enzyme with a 40 µL solution containing 1 mM arabinosyl hypoxanthine (ara H) and 1 mM nucleobase (Ade, 6-methoxyguanine, 2-fluoroadenine, 6-chloropurine) in 50 mM MES pH 6.5 at 40 °C and 300 r.p.m. for 5-240 min. Subsequently, the enzyme was inactivated, following the procedure previously described^[48]^, and the production of arabinosyl adenine (ara A, vidarabine) was analyzed and quantified using HPLC. Each determination was performed in triplicate, with a maximum error of less than 5%. One unit of activity was (U) defined as the amount of enzyme (mg) that produces 1 µmol of ara A per min (IU) under the assay conditions.

The identification and quantification of substrates and products were performed by HPLC analysis using an ACE 5-μm C18-PFP 250 mm × 46 mm column (Avantor-ACE) under the following conditions: i) continuous gradient (100-90% 0.1 M triethylammonium acetate and 0-10% acetonitrile) for 15 min, ii) an isocratic elution (90% 0.1 M triethylammonium acetate 0.1 M and 10% acetonitrile) for 7 min. Retention times for the reference nucleobases and nucleosides, hereafter abbreviated following the recommendations of the IUPAC-IUB Commission on Biochemical Nomenclature, were as follows: adenine (Ade): 10.2 min; 2′-deoxyadenosine (dAdo), 15.5 min; guanine (Gua), 8.2 min; 2′-deoxyguanosine (dGuo), 12.8 min; hypoxanthine (Hyp), 7.6 min; 2′-deoxyinosine (dIno), 12.3 min; 6-methoxyguanine (6-MeOGua), 19.7 min; 2-fluoroadenine (2-FAde), 14.0 min; 6-chloropurine (6-ClPur), 16.1 min. Retention times for the reference non-natural compounds were as follows: ara adenine (ara A), 14.2 min; ara hypoxantine (ara H), 11.1 min, ara 6-methoxyguanosine (ara 6-MeOGuo), 17.1 min; ara 2-fluoroadenosine (ara 2-FAdo), 16.9 min; ara 6-chloropurine (ara 6-ClPur), 18.7 min.

### Influence of pH and temperature on *Lm*PDTN56C activity

To determine the optimal operational conditions for *Lm*PDTN56C in vidarabine synthesis, the effect of pH and temperature on enzyme activity was assayed. The optimum pH was determined under standard arabinosyl transferase assay conditions, using sodium citrate (pH 3–5), sodium phosphate (pH 6–8), MES (pH 5.5-7), Tris-HCl (pH 7-9) and sodium borate (pH 8–10) as reaction buffers (50 mM). The optimal temperature was determined by performing the standard assay across a temperature range of 20-70 °C.

### Thermal stability

Thermodynamic stability was measured by differential scanning fluorimetry in a Rotor Gene™ 6000 (Corbett Life Sciences) essentially as previously described^[31,49,50]^. To this end, 18 µL of a 20 µM solution of *Lm*PDTN56C was mixed with 2 µL 100x SYPRO Orange (Merck kGaA, Darmstadt, Germany). The samples were then subjected to a temperature ramp spanning from 35 to 95 °C at 1 °C/min recording the fluorescent emission (Exc. 460 nm, Em. 510 nm). The melting temperature (*Tm)* was determined as the minimum in the first derivative of the melting curve against the temperature (dRFU/dT *vs*. T), where RFU represents relative fluorescence units. Additionally, kinetic stability was examined by incubating 4 μg of the pure enzyme in 50 mM MES buffer pH 6 for 24 h at different temperatures (40-50 °C).

### Enzymatic synthesis of vidarabine

To assess the potential of *Lm*PDTN56C as a catalyst for the synthesis of arabinosyl nucleosides, the enzyme-mediated synthesis of vidarabine was selected as a model reaction. Operational parameters, specifically substrate concentration and catalyst loading (purified enzyme concentration), were selected for optimization. To this end, 4-12 µg of pure enzyme were added to a 40 µL solution containing 1-10 mM ara H and 1-10 mM Ade in 50 mM MES buffer pH 6. Then, the reaction mixture was then incubated at 40 °C and 300 r.p.m. for 8 h. At periodic intervals, samples were collected, and the arabinosyl transferase activity was evaluated.

### Computational methods

The crystallographic structure of apo *Lm*PDT (PDB ID 6QAI) served as a template to create the complex with dIno or Ara H. Subsequently, 3D models for both *Lm*PDTN56C and *Lm*PDTN56L were built from the aforementioned *Lm*PDT-substrate complexes using PyMol^[51]^. To this end, firstly, the H++ server was used to determine the protonation state of the receptor^[52]^. The geometry of the ligands was optimized using *sqm* methods implemented in the force field toolkit Antechamber before manual docking into the active center using *L. helveticus* PDT-dAdo complex (*Lh*PDT, PDB ID:1S2G) as a template^[53]^,^[54]^. In all simulations, we utilized the *ff19SB* force field for the parametrization of amino acids^[55]^. OPC water molecules were employed as the solvent, extending 12 Å away from any solute atom. Additionally, to maintain the electric neutrality of the system, 8 Na^+^ ions were added. Then, 100 ns restrained molecular dynamics simulations (MD) at 25 °C were conducted using the *pmemd_cuda*.*SPFP* module integrated into Amber22^[56]^ as previously described^,,[6,36,54]^. All dynamic simulations were analyzed using the *cpptraj* module integrated into Amber22.

The analysis of access channels in *Lm*PDTWT, *Lm*PDTN56C, and *Lm*PDTN56L was performed using Caver 3.0.2 software^[57]^. Snapshots from three independent MD trajectories were extracted at 50 ps intervals over 100 ns. Parameters for the probe radius and clustering threshold for identification channels were set to 1.2Å and 3.5Å, respectively. The finalization point of each channel was positioned 3 Å above the substrate position. For the calculation of per-residue free energy decomposition of selected active site residues, we used the MMPBSA.py module integrated into Amber22 (Miller III *et al*., 2012). Visualization of the MDs and channels was accomplished using PyMol^[51]^.

Multiple sequence alignment was carried out using MAFFT-DASH (https://mafft.cbrc.jp/alignment/server/)^[58]^ with the best 961 sequences retrieved using BlastP^[59]^ with the *Lm*PDTWT sequence (Uniprot E9AWJ0) as query. The abundance of each amino acid in the position equivalent to Asn56 was calculated and represented with GraphPad Prism.

## RESULTS

### Improvement of the arabinosyl transferase activity of *Lm*PDT by semi-rational design

The primary advantage of biocatalysts for pharmaceutical syntheses lies in their chemo-, regio-, and stereoselectivity, which frequently enables the simplification of synthetic pathways by protecting group manipulations, resolutions, by-product formation, etc.. Nonetheless, this selectivity can also be a drawback because most enzymes have naturally evolved as “specialists” to work in nature with high affinity for a limited scope of “natural” substrates.

To overcome the limitations of narrow selectivity towards natural substrates, novel enzymes from the microbial genetic diversity can be tailored towards desired properties for a specific reaction^[60]^. To this end, semi-rational approaches, combining stochastic methods of directed evolution with elements of rational enzyme modification overcome specific limitations associated with both directed evolution and rational design approaches^[61]^,^[62]^. In particular, they are well suited to improve properties that can be heavily dependent on active site architecture, such as substrate acceptance or selectivity. ^[42–44]^. Therefore, to maximize the chances for success, we chose to explore the vicinity of the active site^[63]^, resulting in a reduced sequence space ^[35]^.

Positions for saturation mutagenesis were selected within a 5 Å radius of the active site in the crystallographic structure of *Lm*PDTWT according to previous reports of their influence on substrate selectivity from docking studies of the native substrate^[48]^. The active site architecture of *Lm*PDT displays the typical conserved active site for NDTs, formed by amino acid residues from two adjacent subunits (Tyr7, Asn56, Asp79, and Glu85 from one subunit, and Glu121# and Asn128# from the other)^[53][64]^. Within this active site, two distinct pockets can be identified: the 2’-deoxyribose binding site and the nucleobase binding site, as described in earlier studies. On the one hand, Asp79, Asn128#, Glu85 (a catalytic residue positioned by Tyr8), form a network of hydrogen bonds that facilitate the recognition of the 2′ -deoxyribosyl moiety. On the other hand, the nucleobase is recognized through the involvement of side chains of Asn56 (interacting with N9) and Glu121#. Additionally, recent findings reveal an interplay between the catalytic Glu85 and the spatially close Tyr8 and Asn56 residues in trypanosomatid NDTs^[33]^. Then, single-site saturation libraries were generated on Tyr8, Phe15, Asp43, Asn 56, Asp79, Thr82, Glu121, Asn128, Met130 (**Figure 1**).

**Figure 1.**
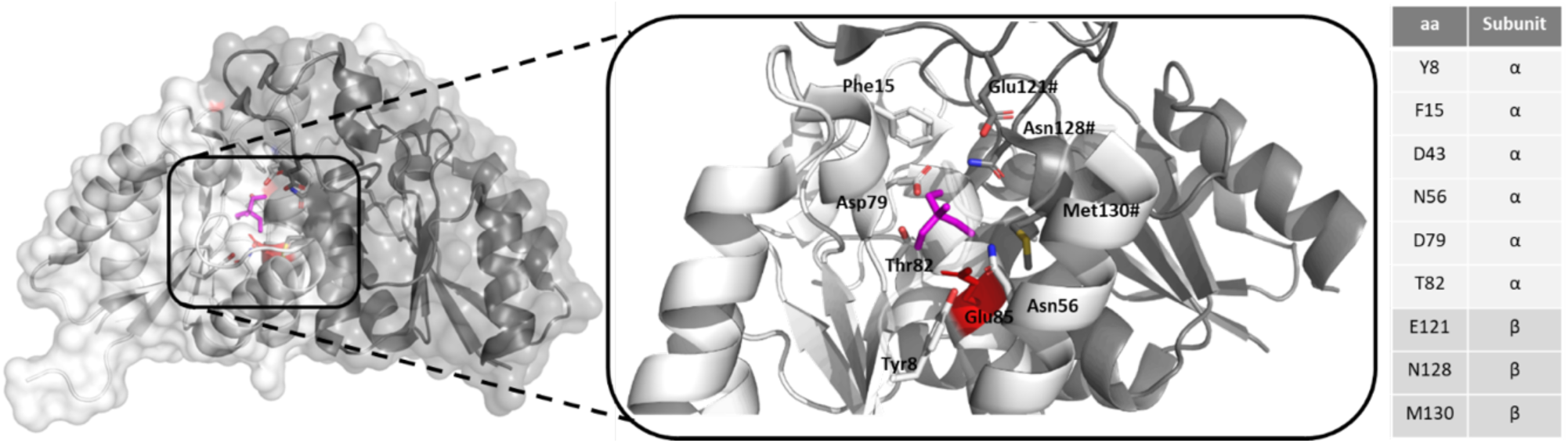
Active site of *Lm*PDT and positions chosen for saturation libraries

Then, a screening or selection method was required to find improved enzyme variants in the libraries. To that end, we leveraged the guanine auxotroph *E. coli* strain PAK6 unable to grow in the presence of guanine nucleoside or nucleoside analogs unless guanine is released by the transglycosylation reaction of NDTs or the phosphorolysis reaction of NPs^[65]^. First, we verified the correct growth in the presence of a known arabinosyl transglycosylase activity using the low-level arabinosyl transferase activity of the parental *Lm*PDT^[48]^. Therefore, we subcloned the gene encoding *Lm*PDTWT with the corresponding RBS and His6 tag-encoding sequences into pBluescriptSK. We verified that the orientation of the cassette placed it under the control of the *lac* promoter present in the pBluescriptSK, necessary for protein expression in *E. coli* PAK6. Then, selection conditions were established, using glycerol as carbon source to avoid catabolic repression of P*lac*. As shown in **Figure 2**, cells harboring the wild-type enzyme grew on selective plates with ara G and Ade after 7 days compared to growth with the natural substrate dG and Ade, which took place in 2-3 days.

**Figure 2.**
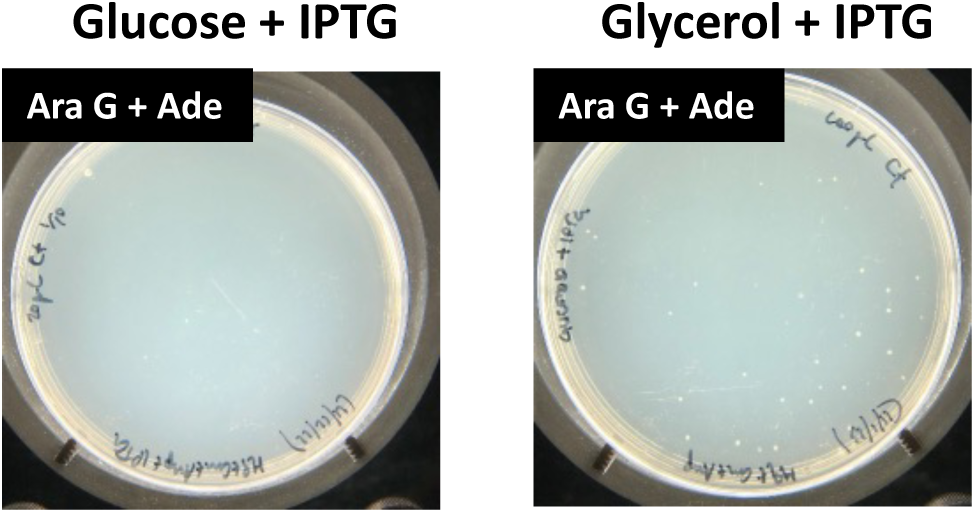
Selection conditions for arabinosyl transferase activity using *E. coli* PAK6 expressing *Lm*PDT_WT_. Images were taken after 7 days of growth.

Next, the nine libraries were transformed into electrocompetent *E. coli* PAK6 as previously reported^[45]^, using the parental *Lm*PDTWT in parallel as a benchmark to detect faster-growing individuals by comparison. While the background arabinosyl transferase activity of the parental *Lm*PDT allowed PAK6 cells to grow within 7 days, 5 colonies grew on the plates corresponding to the Asn56 library within 3-4 days and no other colonies grew in the other 7 libraries. The plasmids of the 5 hit colonies were extracted and sequenced, with 3 colonies presenting the Asn56Leu replacement and 2 presented the Asn56Cys replacement. To rule out phenotypic variability as a cause for the improved fitness, plasmids were subsequently retransformed into *E. coli* DH5⍺ and all five colonies were verified as true hits by HPLC analysis of the arabinosyl transfer reaction from ara H to Ade with crude extracts.

Then, the proficiency of the found *Lm*PDT variants on the synthesis of natural and nucleoside analogues was evaluated with the purified proteins. Interestingly, both *Lm*PDT variants exhibit a substantial increase in activity on vidarabine synthesis, with *Lm*PDTN56L showing up to a 13.5-fold enhancement and *Lm*PDTN56C demonstrating a remarkable 30-fold increase compared to wild-type *Lm*PDT (Table 1). To assess the impact of point mutations on the specificity of *Lm*PDT, both *Lm*PDTN56C and *Lm*PDTN56L were used as catalysts for the synthesis of dAdo from dIno and Ade. As shown in Table 1, the remarkable increase in arabinosyl transferase activity entailed a partial trade-off with 2’-deoxyribosyl transferase activity for *Lm*PDTN56L. However, *Lm*PDTN56C, which suffered a significant reduction of its 2’-deoxyribosyltransferase activity.

**Table 1.** Substrate specificity for *Lm*PDT_WT_ and variants using different nucleosides and nucleobases as substrates.

|  | Donor | Acceptor | Product | Specific activity<br>(IU/mg) |
| --- | --- | --- | --- | --- |
| **2'-deoxyribosyl transferase activity<sup>a</sup>** |  |  |  |  |
| *LmPDT* | | | | 60.7±2.8 |
| *LmPDT*<sub>N56L</sub> | dIno | Ade | dAdo | 47.1±5.2 |
| *LmPDT*<sub>N56C</sub> | | | | 16.2±0.5 |
| **Arabinosyl transferase activity<sup>b</sup>** |  |  |  |  |
| *LmPDT* | | | | 0.004±0.000 |
| *LmPDT*<sub>N56L</sub> | Ara H | Ade | Ara A | 0.054±0.001 |
| *LmPDT*<sub>N56C</sub> | | | | 0.120±0.001 |

<sup>a</sup> Reaction conditions: 0.33 μg of enzyme in 40 $\mu$ l at 40 $^{\circ}$ C for 5 min. Substrate concentrations were 10 mM dIno and 10 mM Ade in 50 mM MES buffer, pH 6.5.
<sup>b</sup> Reaction conditions: 4 μg of enzyme in 40 $\mu$ l at 40 $^{\circ}$ C for 5 min. Substrate concentrations were 1 mM Ara H and 1 mM Ade in 50 mM MES buffer, pH 6.5.

Finally, we attempted to iterate the saturation of the remaining 7 positions on variant *Lm*PDTN56C. However, no colonies grew on the plates with the NNK saturation libraries significantly earlier than in the selection plates with the parental *Lm*PDTN56C. This prevented us from applying an iterative approach which may have introduced mutations already adapted to the arabinosyl transferase enabling substitutions. This lack of additional improvements suggests that protein fitness may have reached a local maximum within the explored sequence space and further advancement will require the exploration of the mutational landscape considering the whole protein structure. Furthermore, the fact that *Lm*PDTN56L has not fully exchanged the deoxynucleotidyl transferase activity for the arabinosyltransferase hints at a “generalistic” nature of this variant, constituting an interesting starting point for further engineering^[66]^. However, a library of 30000 individuals created by error-prone PCR using *Lm*PDTWT as parental and transformed into *E. coli* PAK6 did not yield any hits (data not shown), which suggests that future attempts may require also the exploration of a larger sequence space.

Nevertheless, with a single substitution we achieved variants of *Lm*PDT, with a 30-fold activity improvement towards arabinosyl nucleosides (up to 0.12 IU/mg) with a single amino acid replacement. These activity levels significantly surpass those reported for other NDTs, including *Trypanosoma brucei* PDT (*Tb*PDT, 0.0077 IU/mg, calculated from reported data)^[29]^, *Chroococcidiopsis thermalis* NDT (*Ct*NDT, 0.004 IU/mg, calculated from reported data)^[7]^, and *Lactobacillus reuteri* NDT (*Lr*NDT, 0.0005 IU/mg)^[67]^, making the *Lm*PDT a springboard for further engineering campaigns, addressed at the whole scaffold.

### Biochemical characterization of *Lm*PDT variants

Then, we sought to increase the yield of the reaction through exploration of different physicochemical reaction conditions. First, the pH profile revealed that *Lm*PDTN56C displays high activity across pH range 5.0-6.5, with the highest peak observed when incubated in 50 mM MES buffer pH 6.0 (**Figure** 3). In view of the results, the experimental data suggest a strong pH dependence, which is consistent with the reaction mechanism involving the attack of catalytic Glu85 on C1′ accompanied by a proton transfer to N7 of the purine ring from Glu119 (theoretical pKa 7.2, computed by H++ server, http://newbiophysics.cs.vt.edu/H++/)^[33,54]^. This is also congruent with reports in which, the enzymatic activity of other NDTs is affected by the nature of the buffer,^[33,68]^.

**Figure 3.**
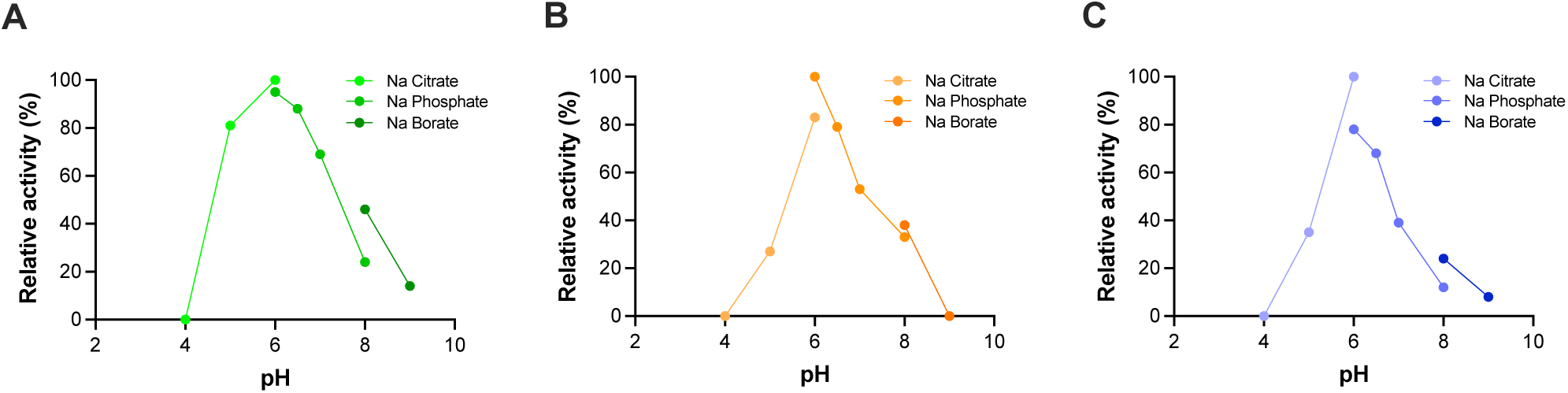
Effect of pH on the arabinosyl transferase activity of (A) *Lm*PDT_WT_, (B) *Lm*PDT_N56L_ and (C) *Lm*PDT_N56C_

Finally, we determined the effect of temperature on the stability and activity of the *Lm*PDT and its variants (**Figure 4**). *Lm*PDTN56C displays high activity, exceeding 60%, within a restricted temperature range of 30 to 40 °C, reaching its maximum at 40 °C, in accordance with previously reported values for the wild-type enzyme^[48]^. The two generated variants were minimally less stable than the wild-type enzyme both at 40 °C and at 45 °C (WT: 56.26 ± 0.10; 55.99 ± 0.04; 55.17 ± 0.24). Regarding their thermodynamic stability, the melting temperatures of the two variants mutant were very only slightly lower than that of the wild-type enzyme (WT: 56.26 ± 0.10; 55.99 ± 0.04; 55.17 ± 0.24) leading us to conclude that the increase in arabinosyl transferase activity did reduce, but not compromise, protein stability. In agreement to other works where focused directed evolution was employed^[39,69]^, the divergence of protein sequences from those more abundant in the natural diversity may be linked to stability losses. Therefore, we explored the natural abundance of the N56C and N56L replacements among NDT sequences. To that end, 10000 candidate NDTs were recovered by BlastP with *Lm*PDT as query and filtered to exclude those that did not contain Asn56, resulting in a set of 9405 sequences. These remaining sequences were subjected to multiple sequence alignment and upon analysis, the abundance of Cys (4.6%) was ten times lower than that of Asp (48.7%) and Asn (44.1%) and leucine was only present in 0.2% of the sequence dataset (Supplementary Figure 1).

**Figure 4.**
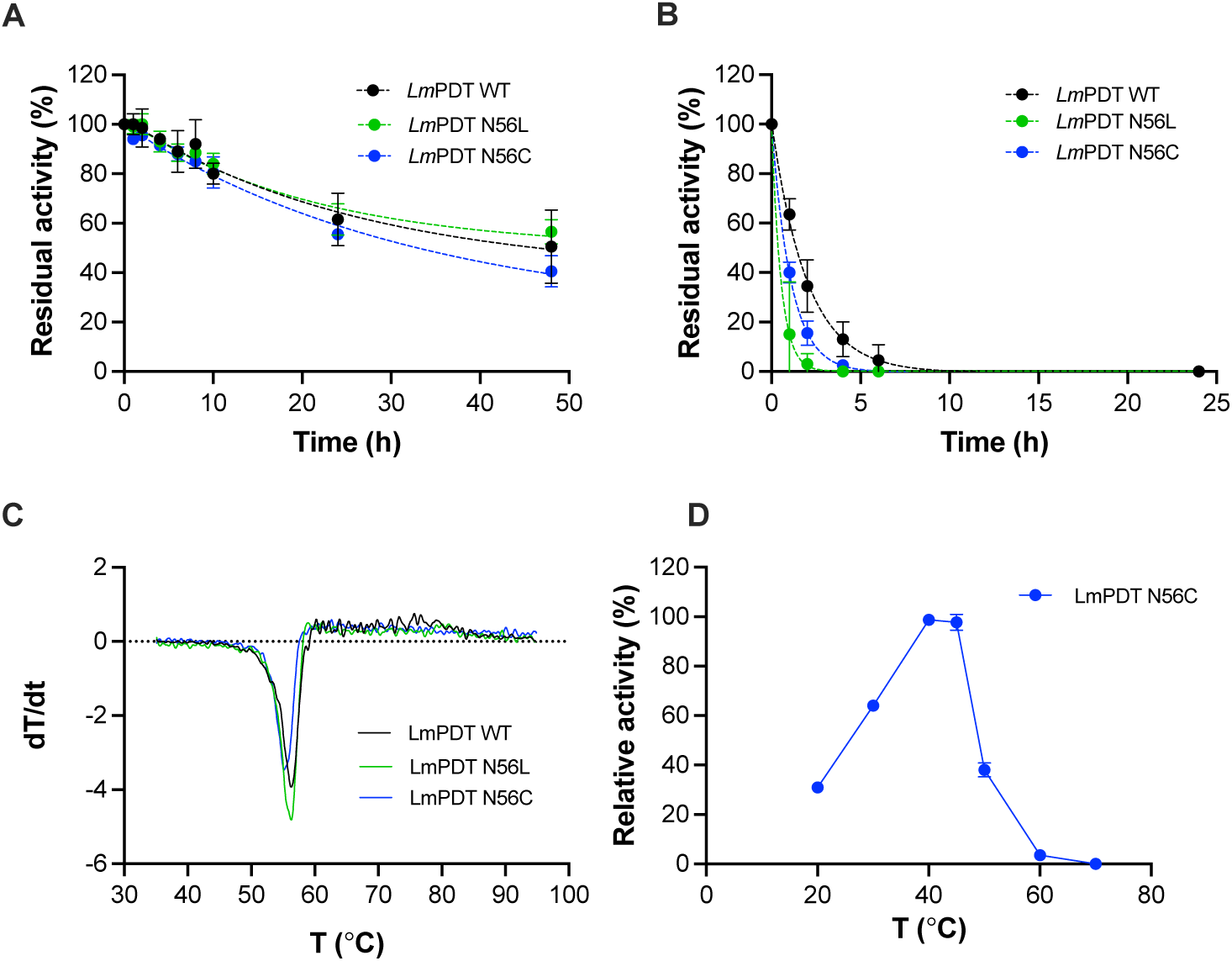
Effect of temperature on the activity and stability of the LmPDT and its N56L/C variants. (A) Thermal inactivation course of *Lm*PDT_N56C_ at 40 °C and (B) 45 °C, (C) first derivative of the melting curves; and (D) optimum temperature for the arabinosyl transferase activity of the N56C variant. Thermal inactivation courses were adjusted to a one-phase exponential decay model (discontinuous line, n=2). Melting temperatures were calculated as the average of n=3 independent determinations.

Finally, as a result of the biochemical characterization, the vidarabine synthesis was carried out under the optimal operational conditions resulting in an activity of 0.16 IU/mg. This represents a remarkable 40-fold improvement in arabinosyl transferase activity compared to *Lm*PDTWT.

### Enzymatic synthesis of arabinosyl nucleosides

We assayed the ability of the *Lm*PDT variants to transfer the arabinosyl moiety from ara H to additional nucleobase acceptors (Figure 5). We observed that the improvement of the N57L/C variants over the wild-type enzyme in arabinosyl transferase activity was independent of the acceptor nucleobase. This is expected for NDTs, which have a lax selectivity towards the acceptor ^[70]^ but also confirms that the biological selection of the libraries was not biased by the acceptor used (Ade). The percentages of synthesis of achieved ranged between ca. 41% when the acceptor was 6-chloropurine to 52% for the synthesis of vidarabine. These rates are significantly higher than those obtained with the *Trypanosoma brucei* PDT using natural nucleosides or nucleoside analogues as donors^[71]^ and 10-20% lower than those obtained with the *Lactobacillus delbrueckii* NDT, however, using dIno as donor.

**Figure 5.**
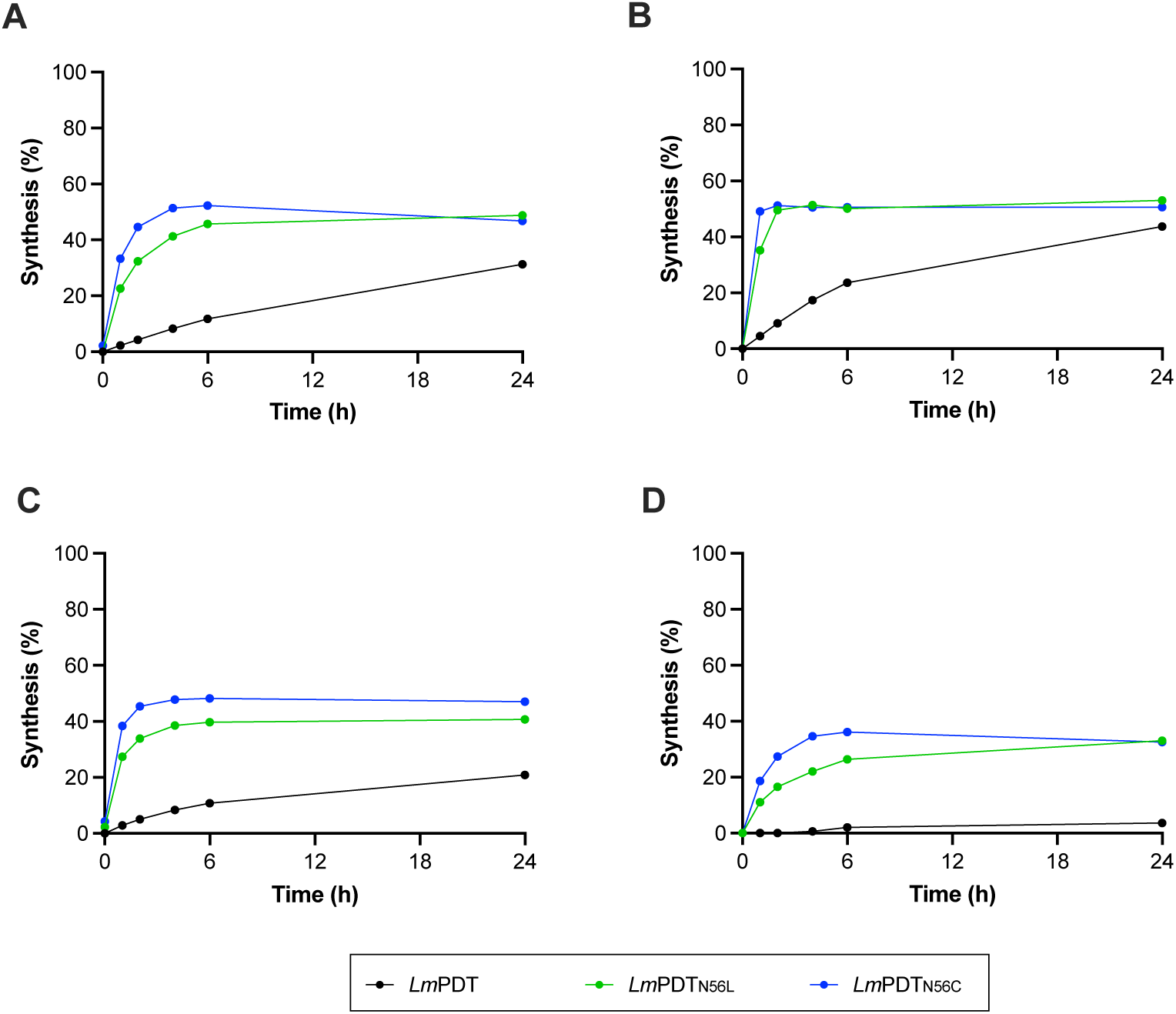
Time-course of arabinosyl transfer activity to of (A) adenine, (B) 6-methoxyguanine, (C) 2-fluoroadenine, (D) 6-chloropurine catalyzed by wild-type and N57L/C variants of *Lm*PDT

Considering that the maximum synthesis was obtained in the synthesis of vidarabine, we chose this reaction as case study, to further probe the influence of substrate concentration and catalyst loading under optimal reaction conditions (Figure 6). We found that *Lm*PDTN56C maintained approximately 50 % conversion when using substrate concentrations ranging from 1 to 10 mM (fixing enzyme concentration), with the maximum µmol converted observed at 10 mM. Furthermore, when holding the substrate concentration constant at 10 mM and varying the enzyme amount, the optimal conversion (%) was achieved with 12 µg of *Lm*PDTN56C.

**Figure 6.**
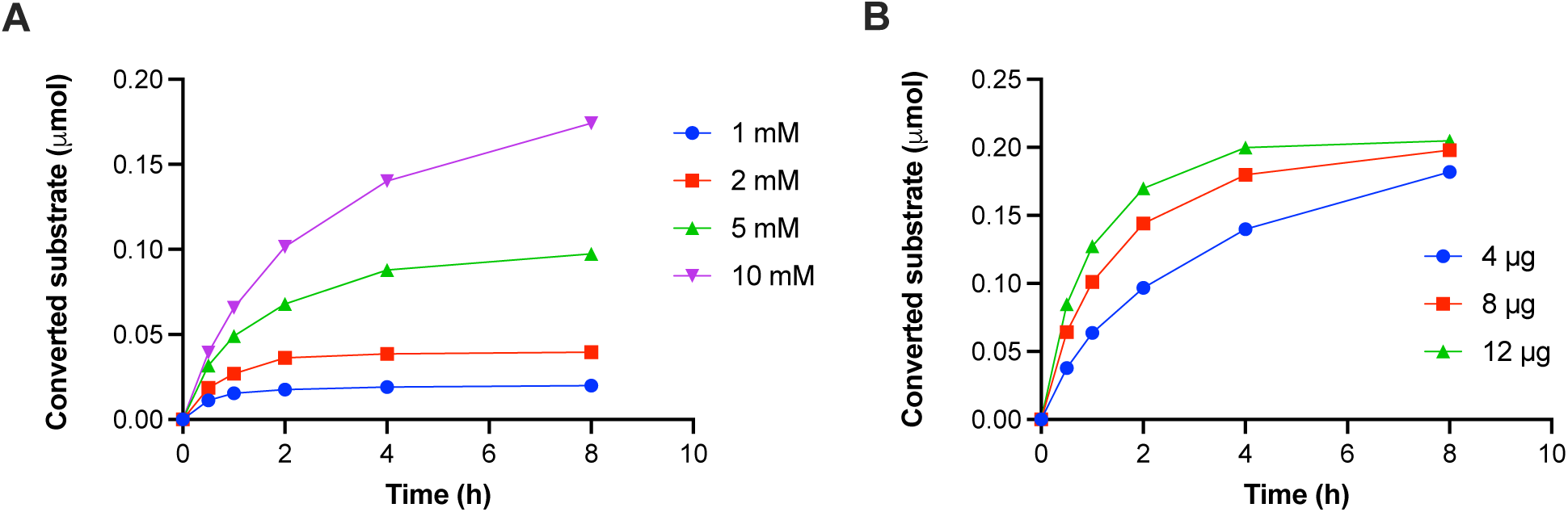
Time-course of enzymatic production of vidarabine catalyzed by *Lm*PDT_N56C_. (A) Influence of substrate concentration. (B) Effect of catalyst loading.

### Determinants of the enhanced arabinosyl transferase activity of *Lm*PDTN56C and *Lm*PDTN56L

To explain the role of the different point mutations on the specificity of *Lm*PDT, 3D homology models of the *Lm*PDTN56L, and *Lm*PDTN56C variants and molecular dynamics simulations were carried out. After a thorough analysis of the initial MD results, no difference in the binding energies of both ligands, dIno and Ara H, within the active site of *Lm*PDT, *Lm*PDTN56L, and *Lm*PDTN56C was found (data not shown). However, following a computational analysis of ligand transport processes from the outside environment into the active site by using CAVER 3.0 software^[57]^ different tunnels and channels were calculated. Based on the throughput score, tunnel 1 was chosen as the most likely entry pathway. Subsequently, we estimated the binding energy of the nucleoside along the principal tunnel. To achieve this objective, CAVERDOCK^[72]^ was employed to calculate the variations in ligand binding energies (dIno or ara H) within both the active site and the surface (τι*EBS*) for the wild-type *Lm*PDT and its respective variants. As shown in Table 2, significant differences in τι*EBS* between dIno and Ara H were found in all cases. These findings indicate a more favorable entry for dIno, consistent with the activity values displayed in in Table 1.

**Table 2.**
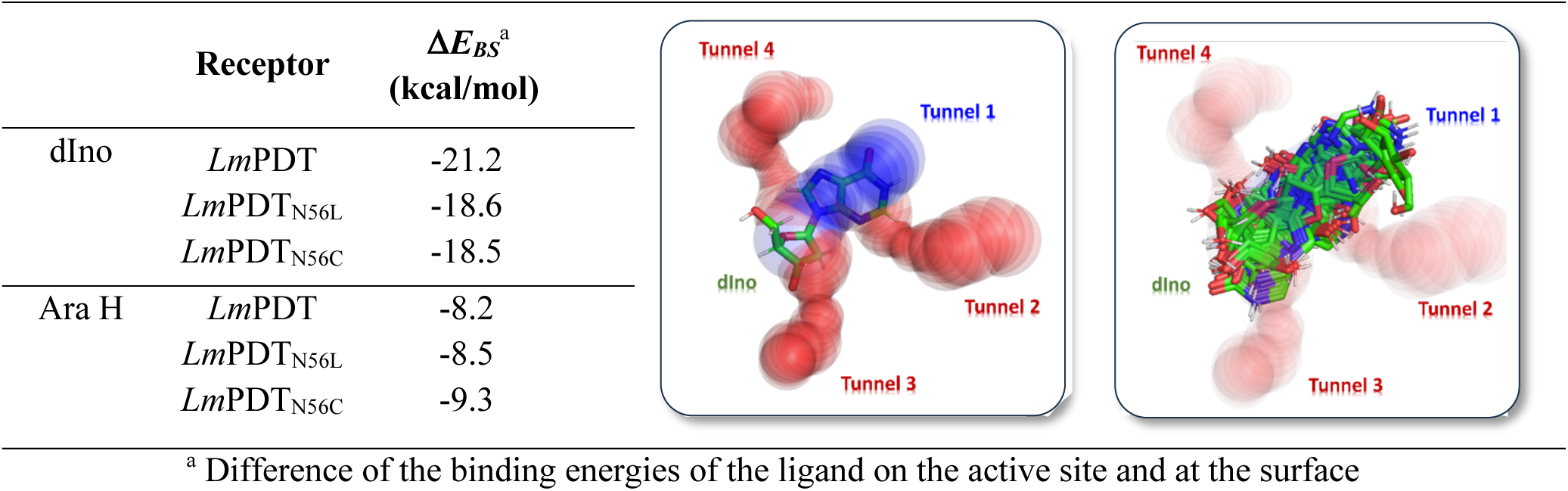
Analysis of ligand transport process in the different receptors along tunnel 1.

Molecular dynamics (MD) simulations conducted on enzyme-ara H complexes for wild-type and variant enzymes revealed different hydrogen-bonding patterns within the active site. As depicted in Figure 2, MD simulations consistently demonstrated an intra-molecular hydrogen bond between the C2’OH and N3 of the purine base. Moreover, as previously described^[48]^, an additional hydrogen bond is observed between N3 and the side chain of Asn56 residue. Notably, Asn56 further forms a hydrogen bond with the OE2 (carboxylate oxygen) from catalytic Glu. Interestingly, the single replacement of Asn56 by Leu (*Lm*PDTN56L) or Cys (*Lm*PDTN56C) led to the disruption of this interplay among Asn56 with catalytic Glu and N3 from the nucleobase (Figure 7). According to experimental findings observed in Table 1, these subtle hydrogen-bonding rearrangements in active site residues seem to tip the balance towards catalysis or the stabilization of a nonproductive complex. To further investigate the arabinosyl transferase activity gain of *Lm*PDTN56L and *Lm*PDTN56C (Table 1), we calculated the per-residue free energy decomposition of the active site residues by MM-PB(G)SA method^[73]^. As shown in Table 2, substitutions N56L and N56C lead to a reduction in the interaction energy within the residue 56-ligand and residue 56-catalytic Glu pair interactions. This reduction may lead to the rearrangement of ara H into the active site, facilitating the stabilization of a productive complex, which in turn, contributes to an improved catalytic process.

**Figure 7.**
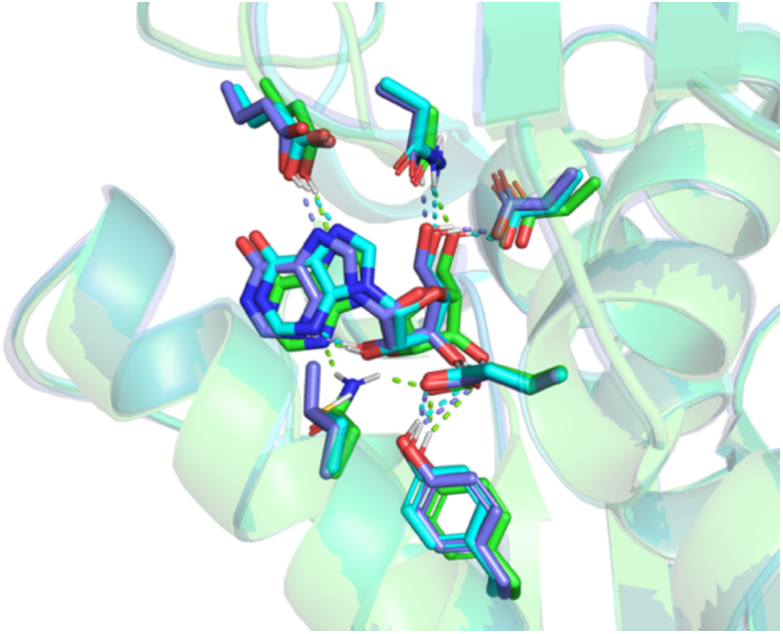
Close view of ara H lodged in the near-attack conformation within the active sites of *Lm*PDT (green carbon), *Lm*PDTN56L (dark blue carbon), and *Lm*PDTN56C (cyan carbon). Hydrogen bonds are represented as colored dotted lines. The figure was prepared with PyMOL (http://www.pymol.org/pymol)

Similarly to other NDTs, the catalytic mechanism of *Lm*PDT involves two ‘half-reactions’ and a common covalent enzyme intermediate. The first half-reaction involves glycosidic bond cleavage by nucleophilic attack of Glu85 on C1′ accompanied by proton transfer from Glu121# to N7 of the purine ring. Furthermore, a second nucleobase occupies the place of the released nucleobase, and the second half-reaction (transglycosylation) follows a similar mechanism but in the reverse direction,^[19,54]^.

This is in accordance with a ping-pong bi-bi mechanism, the established kinetic pathway for NDTs. The catalytic efficiency in this mechanism relies on the optimal availability and access of both substrates to the active site. If the initial substrate (nucleoside) encounters hindrances in accessing the active site, it may lead to a diminished reaction rate. Modulating the access to tunnels and active sites of enzymes has been described before either deliberately pursued as part of a rational design strategy or as a serendipitous solution found by directed evolution. For instance, modulating the access tunnel in enzymes, such as hydrolases or P450 monoxygenases has been described to create selectivity towards shorter chain length,^[74]^, insaturations^[75]^ or different enantiomers^[76]^.

As expected by the previous results, the enzyme-mediated synthesis of vidarabine catalyzed by *Lm*PDTN56C considerably improved previous attempts reported for other NDTs, commonly associated with suboptimal conversions, prolonged reaction times (24-48 hours) and low substrate concentrations (1 mM) ^[29,36,37,67]^. This can be extended to other vidarabine synthesis processes catalyzed by different enzyme families, like nucleoside phosphorylases^[17]^. Despite the successful implementation of several examples of one-pot, two-step vidarabine synthesis catalyzed by a combination of nucleoside phosphorylases (uridine phosphorylase, UP, and purine nucleoside phosphorylase, PNP), yielding moderate to high conversion rates at medium-high substrate concentrations (5-10 mM), long reaction times are needed to reach maximum conversion (24-48 h)^[77,78]^. In this context, the one-pot, one-step approach provided in this work offers a similar operational frame but boasts shorter reaction times (4 hours). Additionally, given the pivotal requirement for catalyst reusability in industrial applications, the immobilization of *Lm*PDTN56C allows the development of a homogeneous catalyst with controlled stability. This stands in contrast to the necessary co-immobilization of both UP and PNP, leading to a heterogeneous catalyst with different stabilities. This, in turn, contributes to increased overall production costs due to the complexities associated with maintaining the catalyst’s performance.

## CONCLUSIONS

This study addresses the inherent limitations of 2’-deoxyribosyltransferases (NDTs) in their catalytic scope, specifically their restricted activity towards substrates with modifications at the 2′-C and 3′-C positions of the ribose moiety. Through a strategic structure-based directed evolution approach, we successfully developed two engineered variants of *Lm*PDT, *Lm*PDTN56L and *Lm*PDTN56C, which exhibit 13.5- and 30-fold enhanced activity in vidarabine synthesis, respectively. The thorough biochemical characterization, encompassing pH and temperature ranges, allowed the optimization of the operational conditions. Furthermore, thermal denaturation experiments underscore its resilience, with an apparent unfolding temperature (*Tm*) of 55.1 °C and retained activity of approximately 61% after 24 hours at 40 °C. As a proof of concept, our investigation into increasing substrate concentration and reduction of catalyst loading emphasizes the potential of *Lm*PDTN56C, particularly in the synthesis of vidarabine. This comprehensive study positions the engineered *Lm*PDT variants as promising tools for efficient nucleoside analog production, opening avenues for further engineering towards enhanced applications in pharmaceutical synthesis.

## AUTHOR CONTRIBUTIONS

DC: investigation, methodology, formal analysis, writing – original draft preparation, writing -review and editing; JA: investigation, formal analysis, writing – original draft preparation, LPS: investigation; CO: investigation; PAK: methodology, writing -review and editing; JFL: supervision, writing – original draft preparation, writing -review and editing, funding acquisition; AH: conceptualization, supervision, writing – original draft preparation, writing – review and editing, funding acquisition, project administration.

## ACKNOWLEDGEMENTS

The authors gratefully acknowledge support with multiple sequence alignments from D. Abia of the Bioinformatics core facility at CBMSO.

## SUPPLEMENTARY FIGURES

**Supplementary Figure 1.**
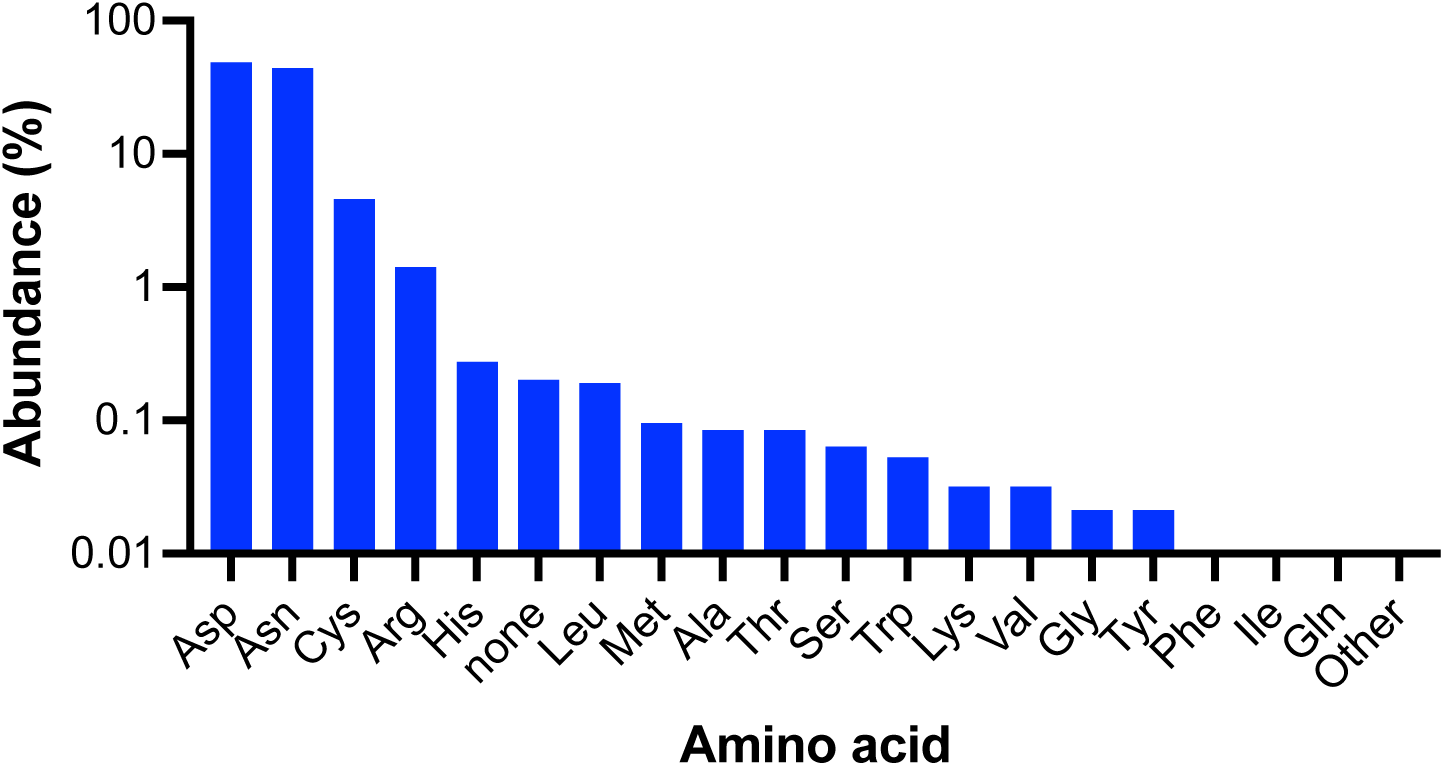
Amino acid distribution of the equivalent position of Asn56 of *Lm*PDT in a set of 9405 sequences with homology to *Lm*PDT.

## SUPPLEMENTARY TABLES

**Supplementary Table 1.** Plasmids used in this work.

| Plasmid | Description | Reference |
| --- | --- | --- |
| pET28b-LmPDT | Vector for LmPDT expression |  |
| pBluescript-LmPDT | Vector for LmPDT library construction and LmPDT expression | This work |

**Supplementary Table 2.**
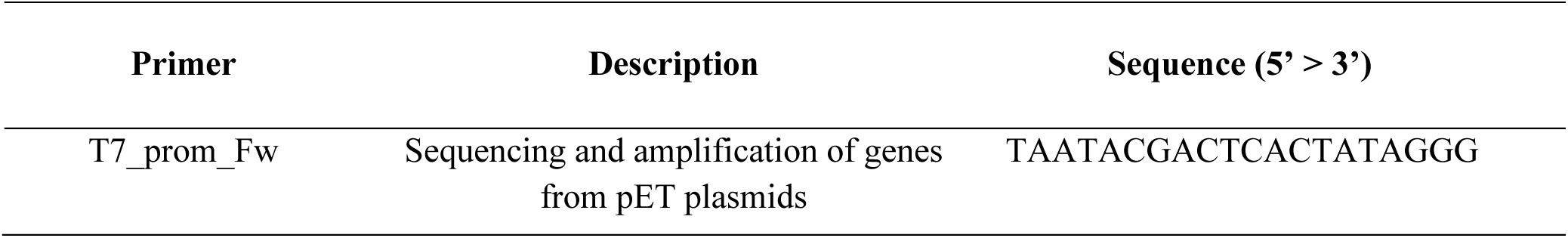
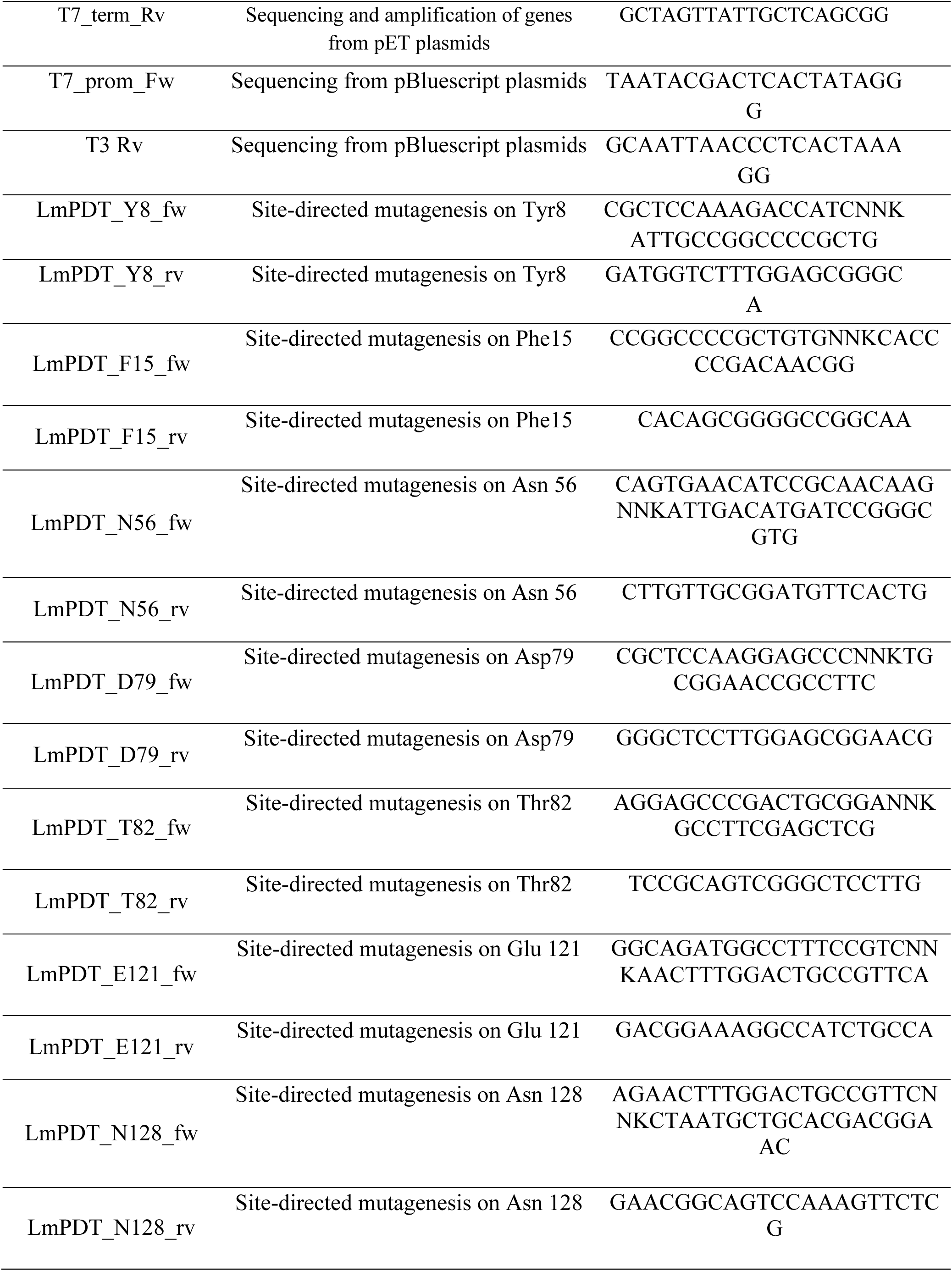
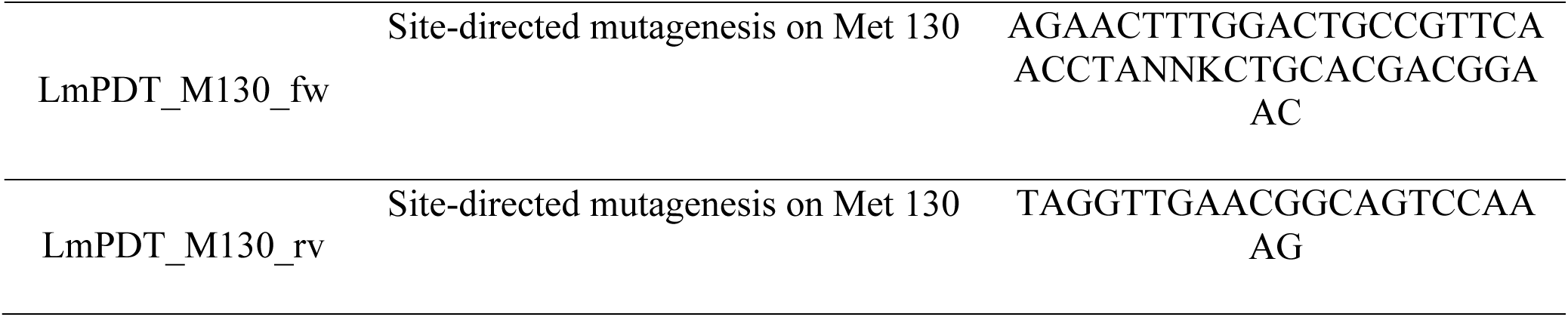
Primers used in this work.

